## Supporting information version3 for "Layer 6 corticocortical neurons are a major route for intra and interhemispheric feedback"

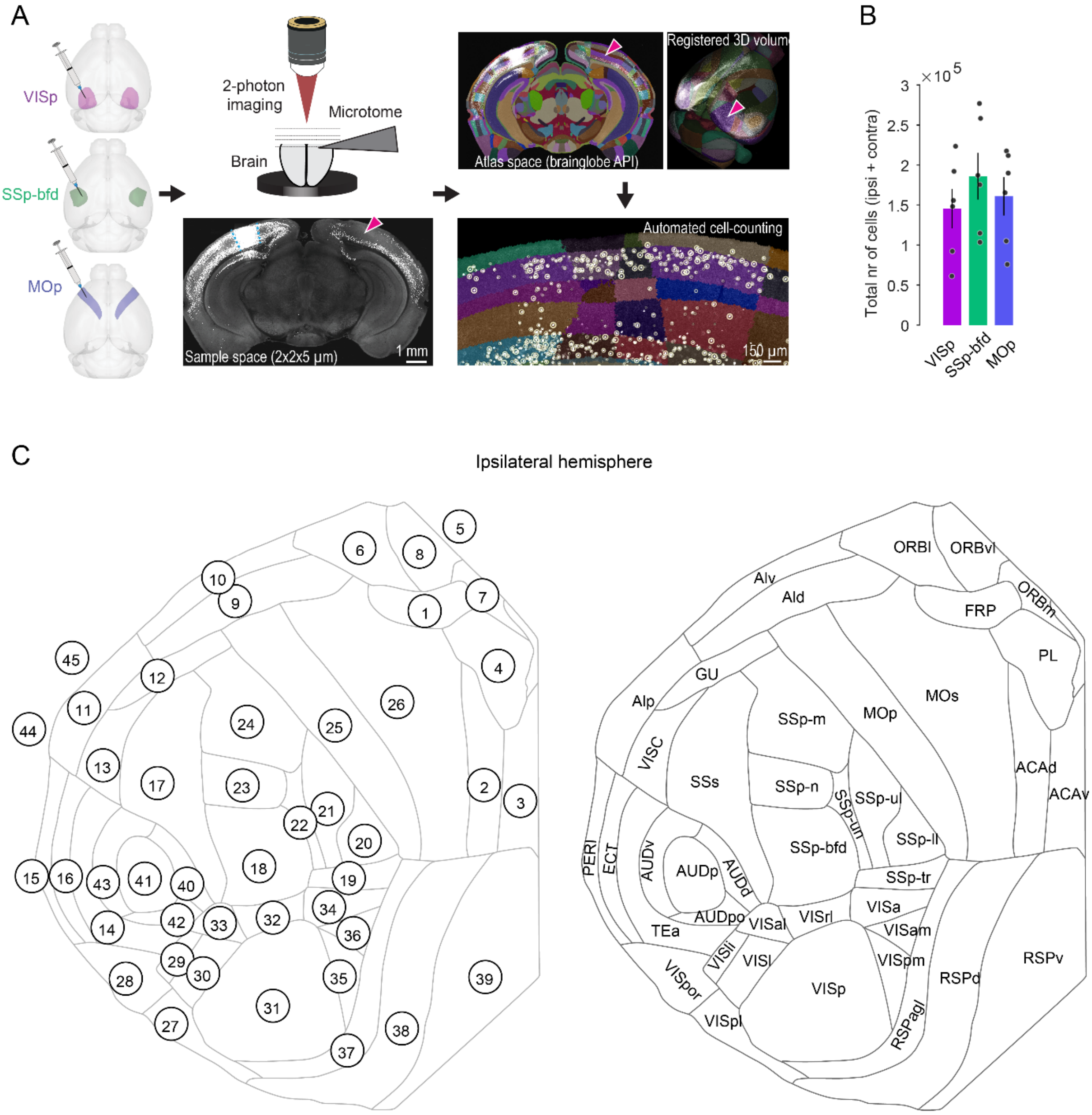


**Fig.S1: Experimental workflow and cell detection.** (A) AAV-EF1a-H2B-EGFP (nuclear retro-AAV) injection into VISp, SSp-bfd and MOp. After viral expression, whole brains are cut and then imaged at x-y-z imaging resolution ~2x2x5 µm using serial 2-photon tomography. Blue dashed line indicates viral injection bolus. Whole brains are mapped onto the common reference atlas (CCFv3) and neurons are automatically detected and counted using the deep-learning based algorithm *cellfinder* (circles). Arrowheads indicate the contralateral counterpart of the injected area (homotopic area) containing retrogradely labeled nuclei of projection neurons. (B) Bar plot displaying average total number of detected cells (± sem) in cortex for the three target areas (n = 6 mice, respectively). (C) Cortical flat map displaying all 45 cortical areas and their identification numbers (left) and abbreviations (right). (1) FRP, frontal pole; (2) ACAd, anterior cingulate area, dorsal part; (3) ACAv, anterior cingulate area, ventral part; (4) PL, prelimbic area; (5) ILA, infralimbic area; (6) ORBl, orbital area, lateral part; (7) ORBm, orbital area, medial part; (8) ORBvl, orbital area, ventrolateral part; (9) ALd, agranular insular area, dorsal part; (10) ALv, agranular insular area, ventral part; (11) ALp, agranular insular area, posterior part; (12) GU, gustatory areas; (13) VISC, visceral area; (14) TEa, temporal association area; (15) PERI, perirhinal area; (16) ECT, ectorhinal area; (17) SSs, supplemental somatosensory area; (18) SSp-bfd, primary somatosensory area, barrel field; (19) SSp-tr, primary somatosensory area, trunk; (20)

SSp-ll, primary somatosensory area, lower limb; (21) SSp-ul, primary somatosensory area, upper limb; (22) SSp-un, primary somatosensory area, unassigned; (23) SSp-n, primary somatosensory area, nose; (24) SSp-m, primary somatosensory area, mouth; (25) MOp, primary motor area; (26) MOs, secondary motor area; (27) VISpl, posterolateral visual area; (28) VISpor, postrhinal area; (29) VISli, laterointermediate visual area; (30) VISl, lateral visual area; (31) VISp, primary visual area; (32) VISrl, rostrolateral visual area; (33) VISal, anterolateral visual area; (34) VISa, anterior visual area; (35) VISpm, posteromedial visual area; (36) VISam, anteromedial visual area; (37) RSPagl, retrosplenial area, lateral agranular part; (38) RSPd, retrosplenial area, dorsal part; (39) RSPv, retrosplenial area, ventral part. (40) AUDd, dorsal auditory area; (41) AUDp, primary auditory area; (42) AUDpo, posterior auditory area; (43) AUDv, ventral auditory area; (44) ENTl, Entorhinal area, lateral part; (45) ENTm, Entorhinal area, medial part. Note that areas (5), (44) and (45) are not represented in the cortical flat map on the right.

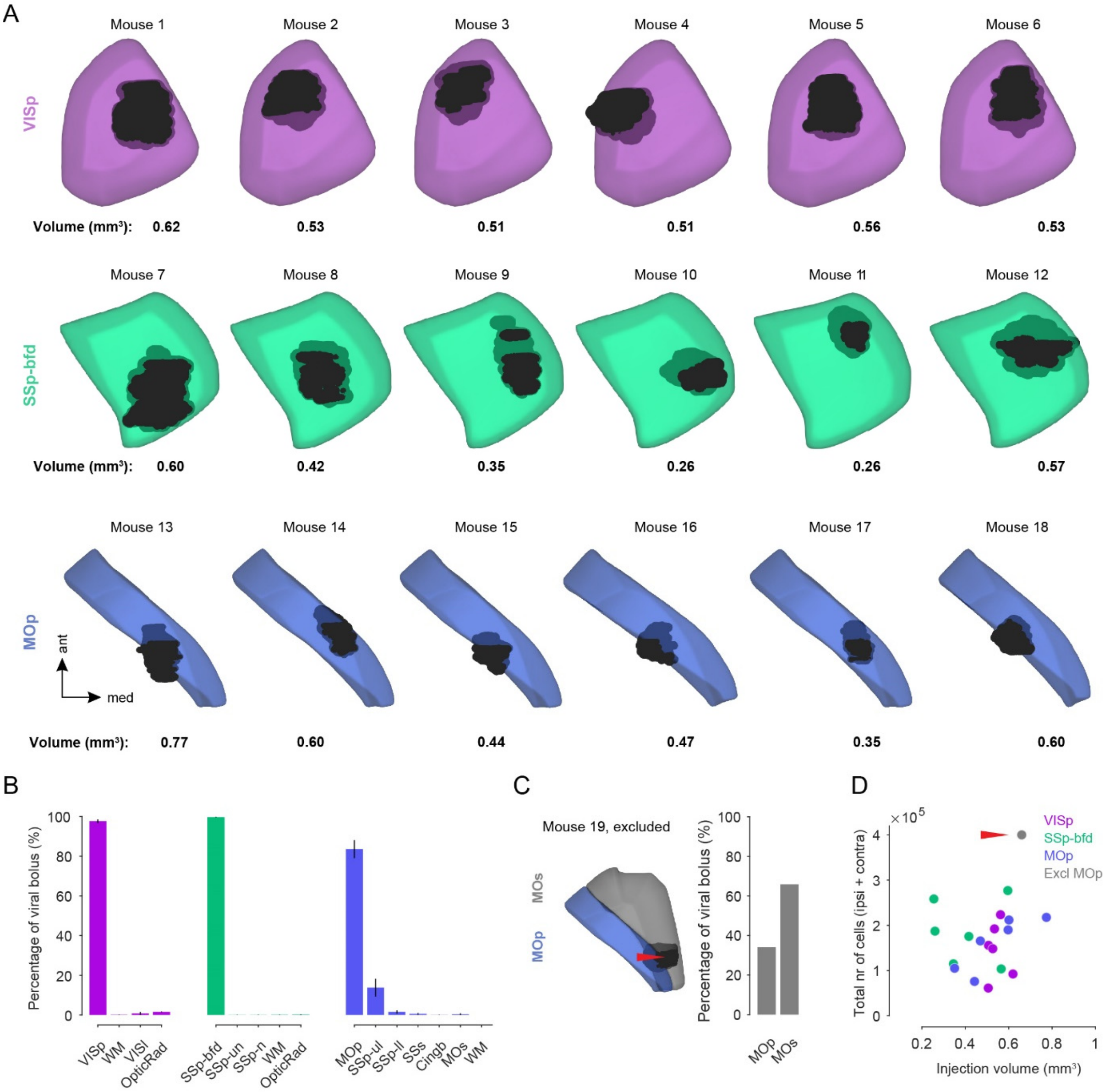


**Fig.S2: Target area viral expression quantification.** (A) 3D reconstruction of the injection site for each mouse warped into the 3D rendered space of VISp (top), SSp-bfd (middle), MOp (bottom) of the CCFv3**.** The value of the volume (in $mm^3$) for each bolus is indicated below. (B) Bar plot displaying the average percentage of the viral bolus (± sem) located in the target and neighboring areas for VISp, SSp-bfd and MOp. WM: white matter) (C) Example of a viral injection targeted in MOp which was excluded from the dataset as the majority of the viral bolus was located in the secondary motor cortex (MOs, red arrowhead) in this mouse. (D) Scatter plot displaying the volume per viral injection versus the total cell number detected in the cortex for each mouse for the three target areas. Red arrowhead highlights injection which was primarily located in MOs and was therefore excluded from the dataset.

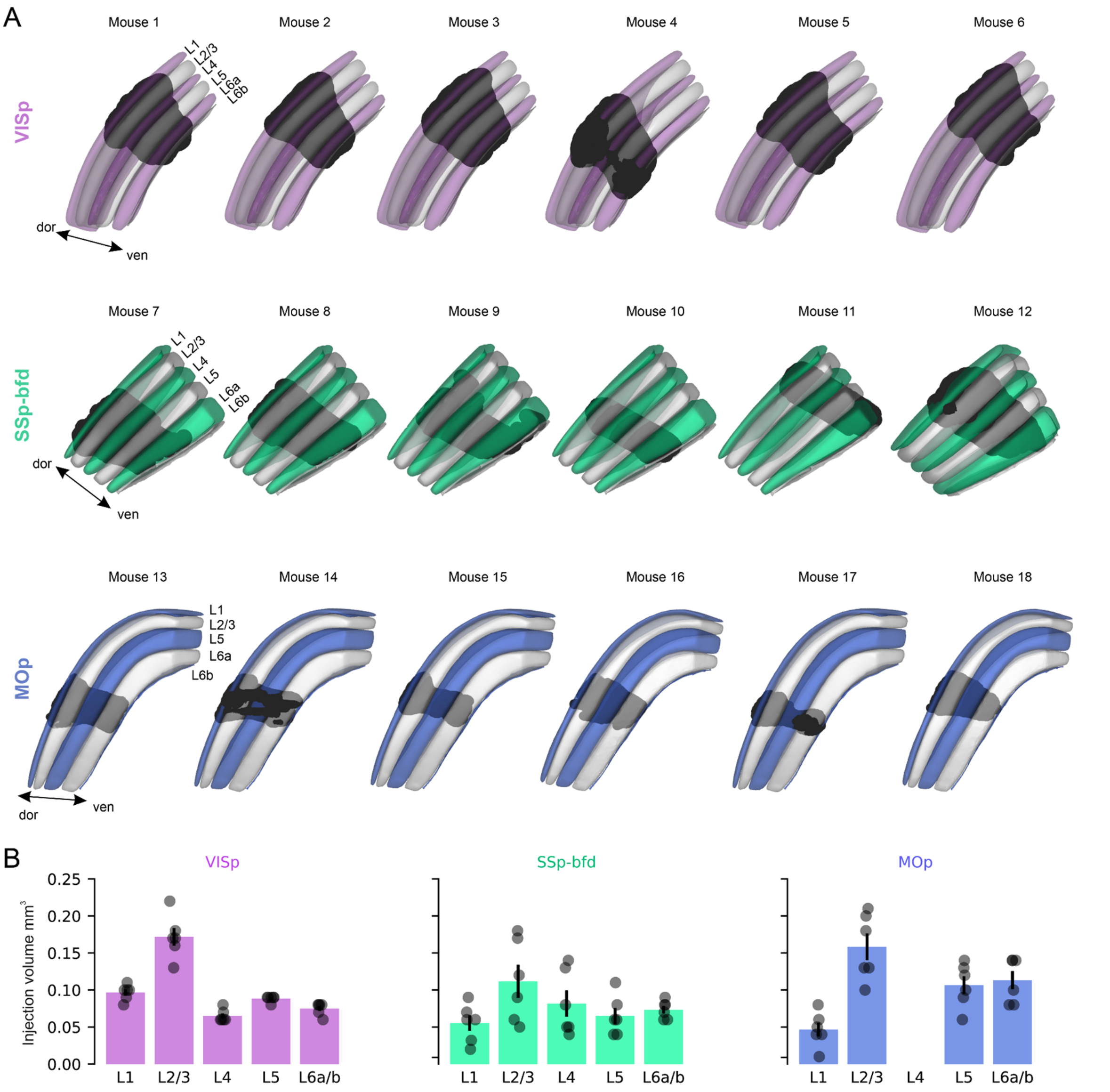


**Fig.S3: Latero-medial view of target area viral expression. (A)** Latero-medial view of the reconstructed injection site for each mouse warped into the 3D rendered space of VISp (top), SSp-bfd (middle), MOp (bottom) of the CCFv3 (see also Fig.S2). (B) Fraction of injection volume per cortical layer for VISp, SSp-bfd and MOp.

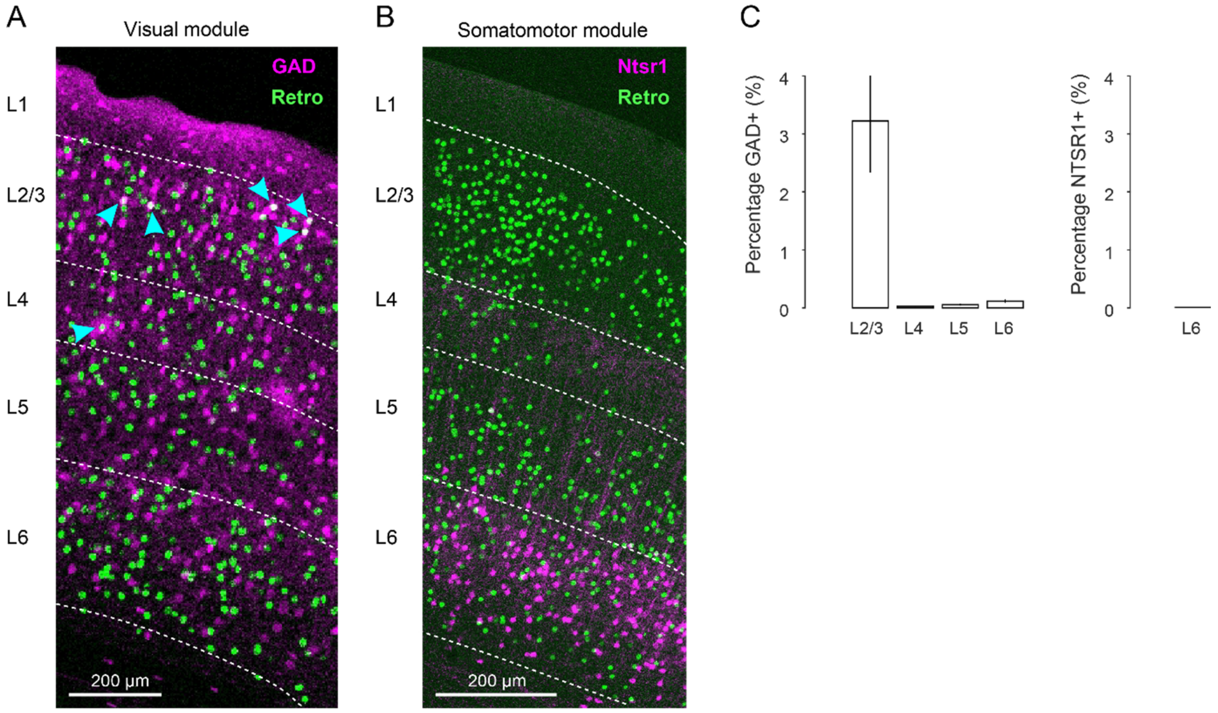


**Fig.S4: Molecular identity of long-range projection neurons.** (A) Confocal image displaying colocalization of GFP+ retrogradely labeled cells and GAD+ (Glutamate decarboxylase) interneurons across cortical layers in an example coronal section of a cortical area of the visual module. Overlap is highlighted with arrowheads. (B) Confocal merged image displaying GFP+ retrogradely labeled cells and NTSR1+ (Neurotensin receptor 1) cells in an example coronal section of a cortical area of the somatomotor module. Note that there is no overlap. (C) (Left) Bar plot displaying the average retrogradely labeled neurons (± sem) that are GAD+ interneurons in L2/3, L4, L5 and L6 across all cortical areas of both hemispheres. (Right) Bar plot displaying the average retrogradely labeled neurons that are NTSR1+ in L6 across all cortical areas of both hemispheres.

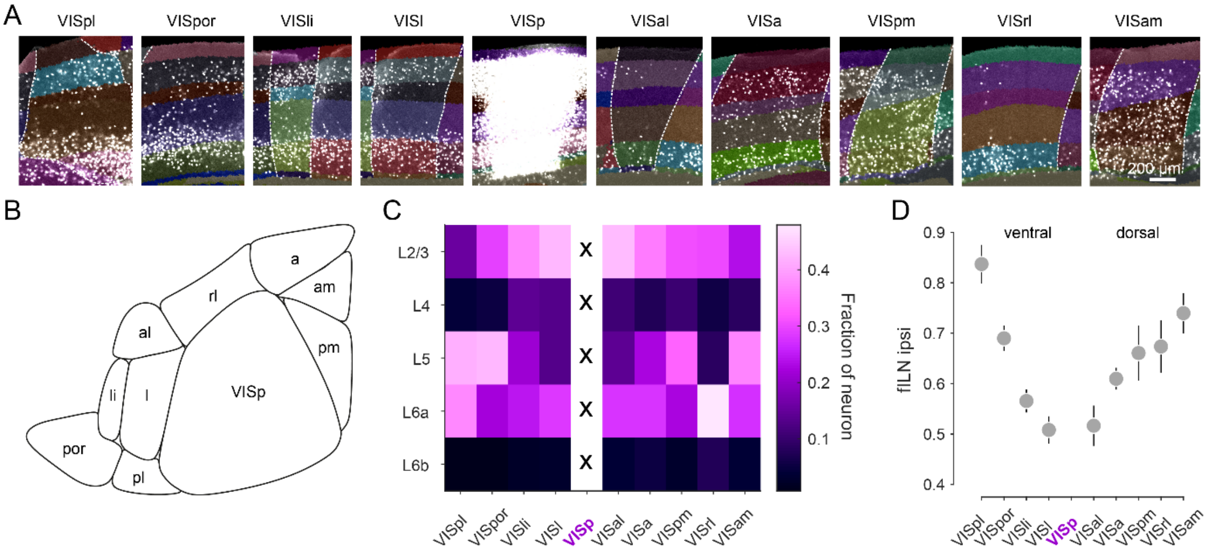


**Fig.S5: Cortical hierarchy of visual areas based on fILN.** (A) Representative images of retrogradely labeled nuclei in higher visual cortical areas in the ipsilateral hemisphere and the corresponding injection site in VISp. (B) Schematic illustration of the anatomical arrangement of VISp with the surrounding higher visual cortical areas in the left hemisphere. (C) Heatmap showing the layer specific average projection weights of higher visual cortical areas onto VISp. (D) Rank ordered average fILN values (± sem) of higher visual cortical areas of the ventral and dorsal visual stream. Injection target highlighted with magenta.

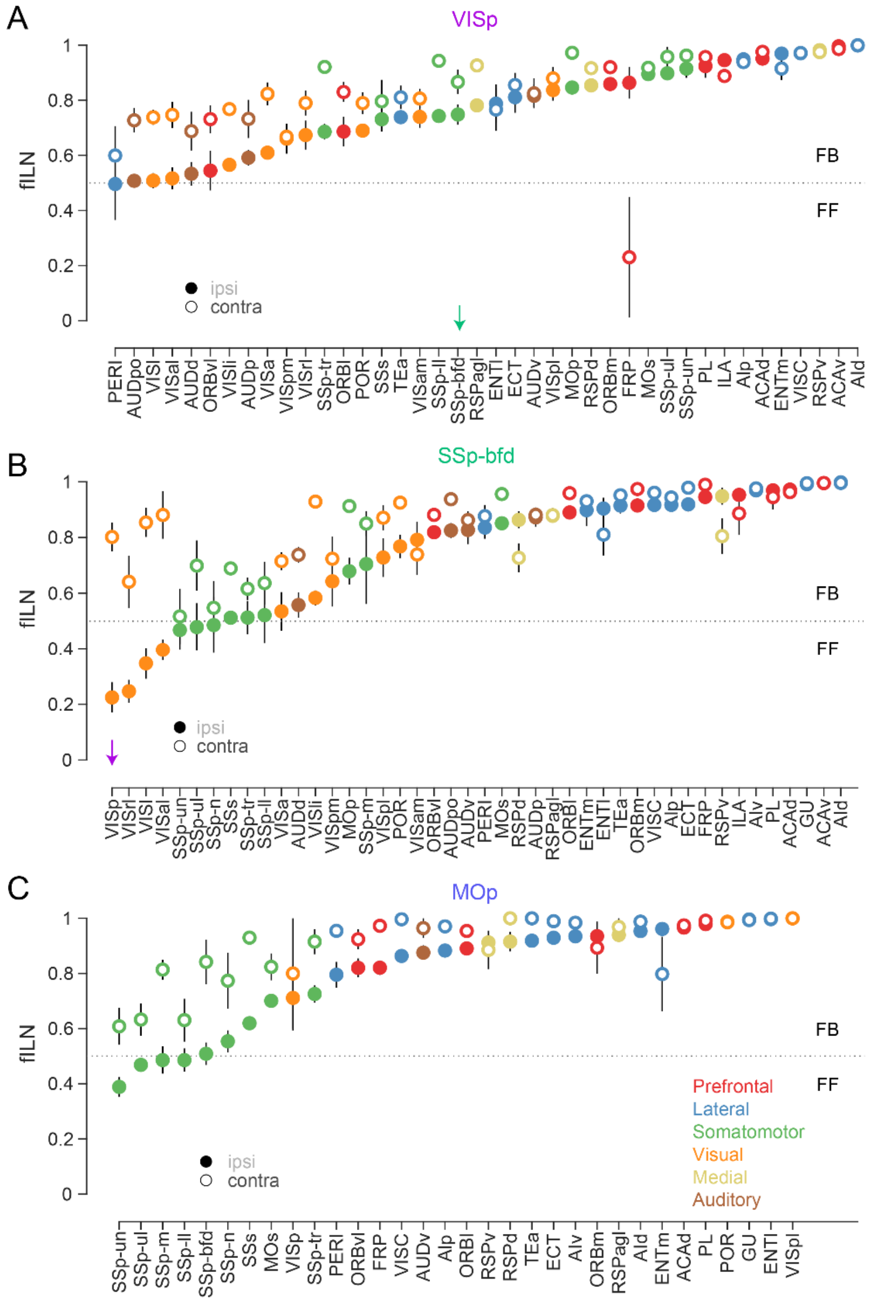


**Fig.S6: Cortical hierarchy per projection area onto VISp, SSp-bfd and MOp.** Plot showing the average ipsilateral (closed circle) and contralateral (open circle) fILN values (± sem) for the individual cortical areas containing both ipsilateral and contralateral projections (excluding the injection target area, respectively) for VISp (A), SSp-bfd (B) and MOp (C). Respective fILN values ranked according to the ipsilateral fILN values. Green arrow in (A) highlights ipsilateral and contralateral feedback projection from SSp-bfd to VISp. Magenta arrow in (B) highlights ipsilateral feedforward but contralateral feedback projection from VISp to SSp-bfd.
